# Surfactant Proteins SP-B and SP-C in Pulmonary Surfactant Monolayers: Physical Properties Controlled by Specific Protein–Lipid Interactions

**DOI:** 10.1101/2022.12.12.520108

**Authors:** Juho Liekkinen, Agnieszka Olżyńska, Lukasz Cwiklik, Jorge Bernardino de la Serna, Ilpo Vattulainen, Matti Javanainen

## Abstract

The lining of the alveoli is covered by pulmonary surfactant, a complex mixture of surface-active lipids and proteins that enables efficient gas exchange between inhaled air and the circulation. Despite decades of advancements in the study of the pulmonary surfactant, the molecular scale behavior of the surfactant and the inherent role of the number of different lipids and proteins in surfactant behavior are not fully understood. The most important proteins in this complex system are the surfactant proteins SP-B and SP-C. Given this, in this work we performed non-equilibrium all-atom molecular dynamics simulations to study the interplay of SP-B and SP-C with multi-component lipid monolayers mimicking the pulmonary surfactant in composition. The simulations were complemented by *z*-scan fluorescence correlation spectroscopy and atomic force microscopy measurements. Our state-of-the-art simulation model reproduces experimental pressure–area isotherms and lateral diffusion coefficients. In agreement with previous research, the inclusion of either SP-B and SP-C increases surface pressure, and our simulations provide a molecular scale explanation for this effect: The proteins display preferential lipid interactions with phosphatidylglycerol, they reside predominantly in the lipid acyl chain region, and they partition into the liquid expanded phase or even induce it in an otherwise packed monolayer. The latter effect is also visible in our atomic force microscopy images. The research done contributes to a better understanding of the roles of specific lipids and proteins in surfactant function, thus helping to develop better synthetic products for surfactant replacement therapy used in the treatment of many fatal lung-related injuries and diseases.

## Introduction

The pulmonary surfactant (PSurf) is a mixture of lipids and surfactant proteins (SPs) that lines the alveoli and thus separates inhaled air from the alveolar fluid. At this interface, PSurf adopts a mono-molecular layer that connects to many types of membrane structures in the aqueous subphase, consisting of either newly-synthesized surfactant or reservoirs of surfactant squeezed out during exhalation.^1–3^ This complex structure facilitates gas exchange between inhaled air and the bloodstream, promotes lung compliance, and prevents alveolar collapse during exhaling.^2,4–6^

The lipid composition of PSurf is fine-tuned to have a melting point very close to the physiological temperature.^4,7,8,8^ This way, the PSurf lipid fraction, the entire PSurf extract, as well as synthetic PSurf mimics display the gel-like liquid-condensed (L_c_) and fluid liquid-expanded (L_e_) phases, as well as their coexistence without the presence of a plateau in the pressure–area isotherm.^9–12^ The L_c_ component allows PSurf to decrease the surface tension of the air–water interface to a very small value and thus prevents alveolar collapse during exhalation, ^13^ whereas the L_e_ component is required to maintain fluidity and allow for rapid spreading of newly-synthesized or squeezed-out PSurf to the interface. ^14^ The central lipid component for the former effect is dipalmitoylphosphatidylcholine (DPPC) with two saturated chains and a high melting point, which allows for its tight packing into the L_c_ phase upon compression at the physiological temperature. ^4^ The other phosphatidylcholines (PCs) with unsaturated chains provide the surfactant with fluidity, whereas anionic phosphatidylglycerol (PG) is responsible for the interactions with SPs.^2,15^ PSurf also contains up to 10 wt% (15 mol%) of cholesterol (CHOL),^4^ whose role in the interfacial monolayer is somewhat unclear,^16^ albeit it promotes the coexistence of liquid-ordered (L_o_) and liquid-disordered (Ld) phases in the interface-attached membrane structures.^17,18^

PSurf also contains four SPs, two of which (hydrophilic SP-A & SP-D) participate in host defense mechanisms.^19^ The hydrophobic pulmonary surfactant proteins (SP-B & SP-C) are essential for the proper mechanical function of the lungs, including rapid surfactant adsorption to the interface and efficient surface tension reduction.^20,21^ Out of these four SPs, SP-B is the most important protein, given that without it, there is no formation of the PSurf monolayer or flow of oxygen to the blood circulation. ^22,23^ The recently resolved supra-molecular assemblies of SP-B are considered to interconnect PSurf layers and thus facilitate the transfer of lipids and gases during breathing cycles. ^15,24,25^ The hydrophobic proteins preferentially partition to the disordered L_d_ phase in lipid bilayers^8,17^ and restrict its dynamics.^26–28^ However, there is no direct experimental data characterizing the phase-preference of SP-B and SP-C in monolayers with coexisting L_e_ and L_c_ phases. SP-B has been demonstrated to affect the structure of the L_c_ phase,^29^ signaling that it might not exclusively locate to the more fluid L_e_ phase. This is corroborated by the fact that the presence of both SP-B and SP-C perturbs the lateral heterogeneity of PSurf by breaking the L_c_ phase into smaller domains.^28,30–34^ This structural change leads to an increase in the L_e_ phase area and thus to an increase in the surface pressure of PSurf monolayers.^33–36^

In addition, the transverse location and orientation of SPs in the surfactant monolayer remain unknown. Studies on lipid membranes have revealed that the inclusion of SP-B has little effect on the acyl chain region,^37,38^ yet it affects the thermodynamic behavior of membranes.^39^ These findings, together with X-ray diffuse scattering (XDS) results,^40^ place SP-B in the head group region in lipid bilayers. XDS experiments^40^ also detect protein density at the membrane core, which is associated with SP-C and agrees with its parallel orientation along the membrane plane. This different positioning of the proteins might explain why SP-B has a larger effect on surface pressure. ^33–36^ Curiously, earlier studies have concluded that SP-C assumes a transmembrane orientation,^41,42^ although there is also evidence suggesting that this orientation is CHOL-dependent. ^43^ Still, it is unclear how these data on SP positioning in a bilayer translate to their positioning and orientation in lipid monolayers at different compression levels, *i.e*. at different steps of the breathing cycle. Furthermore, there is evidence that SP-B homodimers can form higher oligomers in the form of ring-shaped particles with 10 nm diameter,^15,24^ while SP-C has been suggested to form dimers and higher oligomers in PSurf membranes. ^44^ In addition, the oligomeric state of both SP-B and SP-C in PSurf membranes and monolayers can be modulated by SP-B/SP-C interactions. ^45^ Hence, to understand this interplay, the orientation and positioning of SP-B and SP-C complexes in PSurf membranes and monolayers should be resolved.

The effects of lipids and proteins are also tightly-coupled,^22^ and their mutual interactions are relevant for the surface activity, ^46^ lung homeostasis, ^47^ and for the three-dimensional structure of the PSurf. ^2^ However, the specific interactions between proteins and lipids remain somewhat poorly understood. SP-C has demonstrated no lipid preference in some studies^28^ and especially no interactions with CHOL. ^27^ Curiously, some studies observed that SP-C positioning and tilt in bilayers were affected by CHOL, ^43^ which could result from indirect membrane-mediated effects as CHOL in general increases lipid acyl chain ordering. CHOL also inhibits the surface pressure-decreasing ability of PSurf, yet it is restored by the presence of hydrophobic SPs. ^48,49^ SP-B has been demonstrated to interact preferably with PG and CHOL, ^15,28,50^ yet some studies suggest that it prefers PC instead of PG, ^51^ or that it interacts equally little with PC and PG. ^38^ All in all, these discrepancies highlight the need for further studies into lipid–protein interactions in PSurf.

The dynamic conditions and the small scales involved render experimental studies on PSurf challenging. Therefore, despite exhaustive efforts using a plethora of techniques, ^2,22^ several questions still remain unanswered regarding the complex lipid composition and the roles of SPs in providing the lungs with the desired biophysical properties. Notably, it is unclear how PSurf manages to reach a surface tension of a few mN/m, when monolayers *in vitro* consistently collapse at ≈25 mN/m.^52^ In the *squeeze-out mechanism*, L_e_-forming lipids fold into lipid reservoirs in the aqueous subphase, enriching the remaining PSurf in L_c_-forming DPPC. ^53,54^ Alternatively, rapid compression could transform PSurf into a metastable *supercompressed* state that can maintain low surface tensions for extended times without changes in its composition. ^55^ Which mechanism is responsible for lung functioning ultimately comes down to the specific interactions among PSurf lipids and SPs as well as their lateral and three-dimensional organization. Furthermore, understanding lung functioning and the role of proteins and lipids therein is crucial for the development of better synthetic surfactants to treat newborn respiratory distress syndrome without side effects caused by natural extracts. ^56,57^ Finally, understanding how the structures of PSurf are penetrated by pollutants, pathogens,^58^ and surfactant-coated drugs^59^—all of which are able to circumvent our defense mechanisms—has significant implications for human health.

Molecular dynamics simulations have in principle the power to tackle the aforementioned questions.^60^ However, most such simulations^61^ have been performed using coarse grained models^62^ that provide qualitative information at best due to their limited descriptions of the water–air surface tension^62^ and lipid phase behavior. ^63,64^ As we have recently demonstrated, ^65,66^ the former issue also haunts almost all atomistic simulations of lipid monolayers, ^67,68^ and many studies have considered the behavior of SPs in lipid bilayers instead.^40,69,70^ We have recently demonstrated^65,66^ that the combination of the 4-point “Optimal Point Charge” (OPC) water^71^ and the CHARMM36 lipid models^72^ provides an accurate description of lipid behavior at the water–air interface. We also applied this model to multi-component PSurf-mimicking lipid monolayers, for which we successfully reproduced experimentally observed behavior, including an almost quantitative agreement between the surface pressure–area isotherms. ^11^

Here, we extend our previous study (Ref. 11) by incorporating the two hydrophobic SPs (SP-B & SP-C) in our simulations using the compatible and thoroughly tested CHARMM36 protein model. ^73^ The SP-C model is based on a simple alpha-helical peptide with two palmitoylations, ^74^ whereas the monomeric SP-B model is based on recent experimental data and our subsequent refinement. ^15,24^ While in the simulations we focus only on monomeric forms of SP-B and SP-C, experiments can also contain minor fractions of proteins in different oligomeric states. We first validated our model against existing surface pressure–area isotherms and diffusion coefficients measured here using fluorescence correlation spectroscopy. Then, we studied the vertical SP positioning, monolayer heterogeneity, monolayer dynamics, SP phase partitioning preference, and SP–lipid interactions under native nonequilibrium conditions of the PSurf, and compared our findings to the images from our atomic force microscopy imaging.

## Results and Discussion

We performed atomistic MD simulations of 4-component lipid monolayers with the SP-B and SP-C proteins (“SPB” and “SPC” for the corresponding simulation systems, respectively), and in the absence of a protein (“NoP” for “no protein”). The monolayers contained DPPC, POPC, POPG, and CHOL in molar ratios of 60/20/10/10 to mimic the lipid composition of pulmonary surfactant. ^4^ Analogous lipid mixtures were used before in experimental and computational studies and showed to behave similarly to natural extracts. ^11,75–77^ We performed 5 μs-long compression/expansion simulations and included the entire trajectories in the analyses to cover a large range of monolayer compression states, in the area per lipid (APL) range from 90 to 45 Å^2^, corresponding to surface pressures from 0 mN/m up to ≈70 mN/m. Further details of the simulations and experiments as well as their their analyses are provided in the Methods section and in the SI.

### The Simulation Model Reproduces Experimental Behavior

The snapshots from SPB and SPC compression simulations at large (85 Å^2^), intermediate (65 Å^2^), and small (55 Å^2^) APL are shown in Fig. 1. More snapshots are shown in Fig. S1, including the data for the NoP systems. At 85 Å^2^, all monolayers display disordered lipid acyl chains without neither regularly-packed regions or collective tilting, characteristic of the L_e_ phase. At 65 Å^2^, lipids display more ordering, and the movies at DOI:10.6084/m9.figshare.20375745 reveal the emergence of transient L_c_-like clusters in the monolayer. Still, no clear phase-separation is visible in Fig. 1. At 55 Å^2^, the hexagonal packing of lipid chains is immediately obvious, although not all lipid chains seem to participate in it, preventing the monolayer from fully adapting to the L_c_ phase due to the presence of lipids with unsaturated chains. ^11^ Overall, it seems that the SPs have a relatively small effect on the overall monolayer structure. Still, it seems that the lipids in the vicinity of the proteins display somewhat lower ordering and packing.

**Figure 1:**
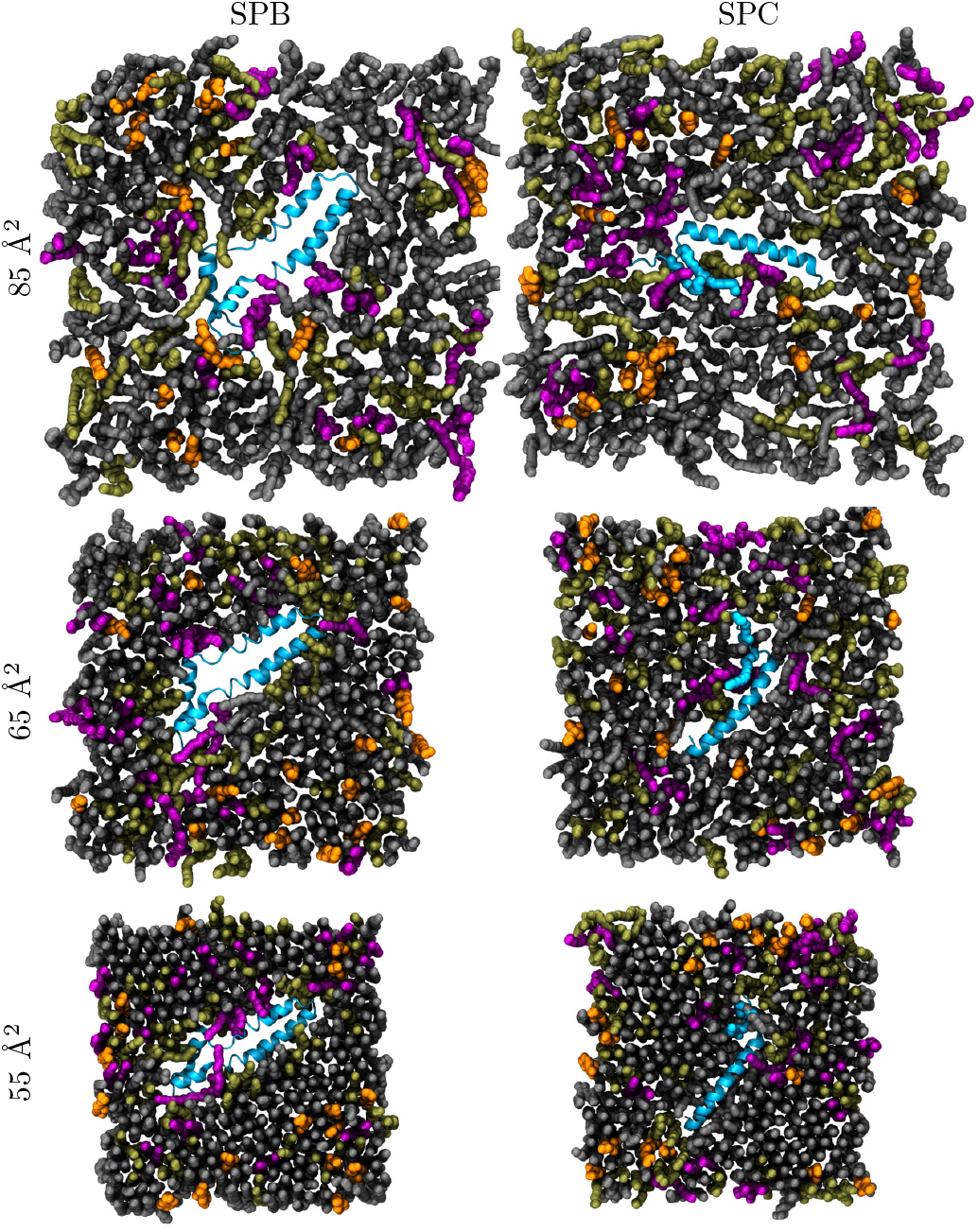
The simulation snapshots. The monolayers are depicted from the air (vacuum) side of the interface. DPPC is shown in gray, POPC in olive green, POPG in purple, and CHOL in orange. Water molecules and hydrogen atoms of lipids are not shown for clarity. The proteins are shown in blue with the ribbon representation.

The SP structures seem stable during the compression simulations with one replica for both SP-B and SP-C proteins temporarily reaching root mean squared deviation (RMSD) of ~1 nm during the simulation (Fig. S2). However, the other replicas show RMSD values of less than 0.5 nm, indicating high stability. The protein structures also become less mobile upon compression, as demonstrated by the RMSD values extracted with respect to the previous protein structure (Fig. S3). This decrease in fluctuations is not surprising considering the increased pressure from the neighbouring lipids, but it also demonstrates that the compression does not lead to some abrupt conformational changes.

Surface pressure–area isotherms of lipid monolayers provide a straightforward way to experimentally characterize surfactant behavior at the air–water interface and to quantify the effect of SPs on monolayer packing. Isotherms are also regularly used to study the binding of proteins or other molecules to lipid membranes, and the changes in surface pressure provide hints on the structural changes caused by the bound molecules, as well as their location. Such an approach, applied earlier to SPs, has revealed that both SP-B and SP-C increase the surface pressure, albeit SP-B has a more significant effect.^33–36^ Surface pressure–area isotherms are also readily extracted from the pressure components which are standard output of MD simulations (see Methods in SI). Due to the non-equilibrium nature of our simulations, both the APL and surface pressure were binned into a histogram using a time window of 100 ns. The isotherms for all compression simulations are shown in Fig. 2. For comparison, we also included an experimental isotherm of a protein-free lipid monolayer with the same composition as the simulated ones from our previous work. ^11^

**Figure 2:**
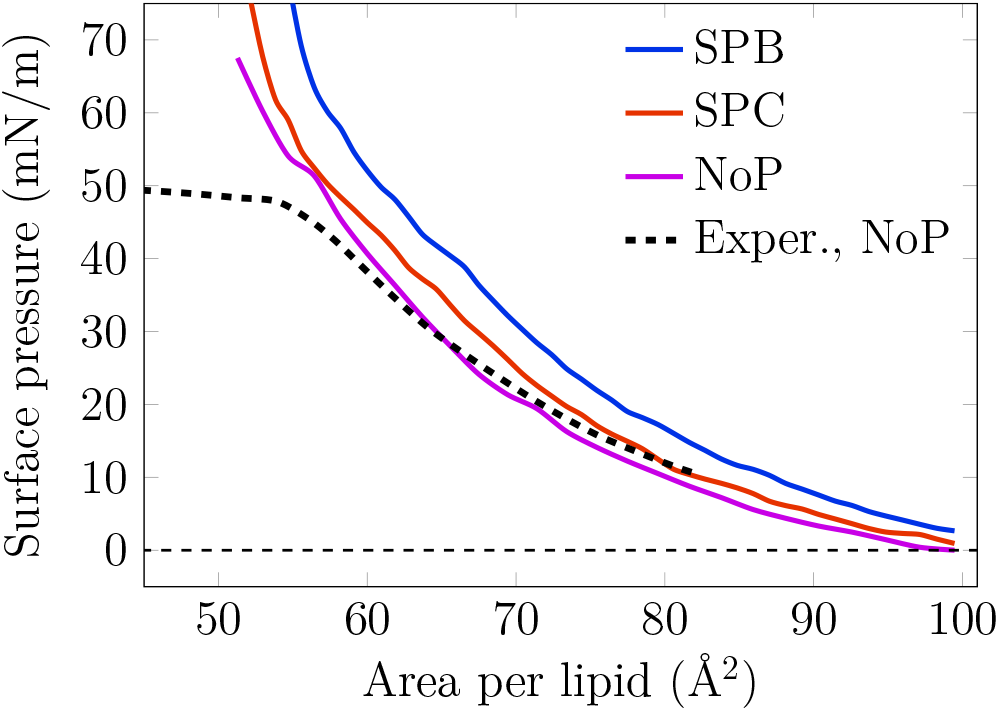
Simulated surface pressure–area isotherm for the monolayer with SP-B or SP-C, and without any proteins at 310 K. The curves are extracted from compression simulations. Experimental compression isotherm data for the protein-free system is taken from Ref. 11. Full data set for the NoP system, including compression and expansion simulations at both 298 K and 310 K, are shown in Fig. S4.

The isotherms for the NoP system agree well between the simulations and experiments. The system without proteins was also subjected to an expansion simulation, in which the area was increased at the same rate. The isotherms from compression and expansion simulations of the NoP systems (see Fig. S4) reveal no significant hysteresis at 310 K. However, there is some hysteresis at 298 K and at the relatively low APL where the L_c_ phase dominates, yet this is not surprising considering the simulation time scale. The APL range corresponding to the L_e_ phase is free of hysteresis effects at both studied temperatures. The size of the simulated monolayer patch in atomistic simulations is too small to allow for the monolayer to properly collapse at a pressure of ~45–50 mN/m that is observed in Langmuir trough experiments.^52^ Still, the agreement between experimental and simulated isotherms is generally excellent. Moreover, the isotherms derived from static equilibrium simulations (in Ref. 11) and dynamic non-equilibrium simulations (this work) are similar (see Fig. S4), signaling that the compression rates available to atomistic simulations are sufficient to reach quasi-equilibrium.

The SPs affect the surface pressure–area isotherm in distinct ways. Here in the SPB system, the presence of SP-B increases the surface pressure across the isotherm by 7.6±1.4 mN/m without changing its shape (system “SPB”). This is in line with the increase observed in Langmuir trough experiments, ^34–36^ and indicates that the presence of SP-B promotes the L_e_ phase by somehow perturbing the tightly-packed L_c_ phase. The role of SP-C on surface pressure based on experiments is less clear, ^33^ yet it has also been suggested to increase surface pressure^36^ albeit less than SP-B. We indeed observe a smaller upward shift of 2.6±1.6 mN/m across the isotherm in the presence of SP-C in our simulations (system “SPC”).

In addition to surface pressure, another property sensitive to compression is the lateral diffusion of the molecules in the monolayer. To further validate our simulation model and to evaluate the effects of proteins on monolayer behavior, we performed *z*-scan fluorescence correlation spectroscopy (FCS) measurements on two monolayers (see Methods for details): a protein-free “NoP” composition, and poractant alfa—an extract of the natural porcine pulmonary surfactant that contains polar lipids and hydrophobic surfactant proteins yet lacks nonpolar lipids. We thus re-introduced 10% of CHOL in the poractant alfa to mimic natural surfactant, following our earlier approach. ^75^ Atto633-labeled DOPC was used as a probe. In simulations, diffusion coefficients were extracted by fitting displacement distributions of lipids over a 10 ns time interval (see Methods in SI for details). These methodologically different approaches should not affect the conclusions, since all components in the same membrane phase are expected to have the same diffusion coefficients.^78^ The diffusion coefficients extracted from simulations and experiments are shown in Fig. 3.

**Figure 3:**
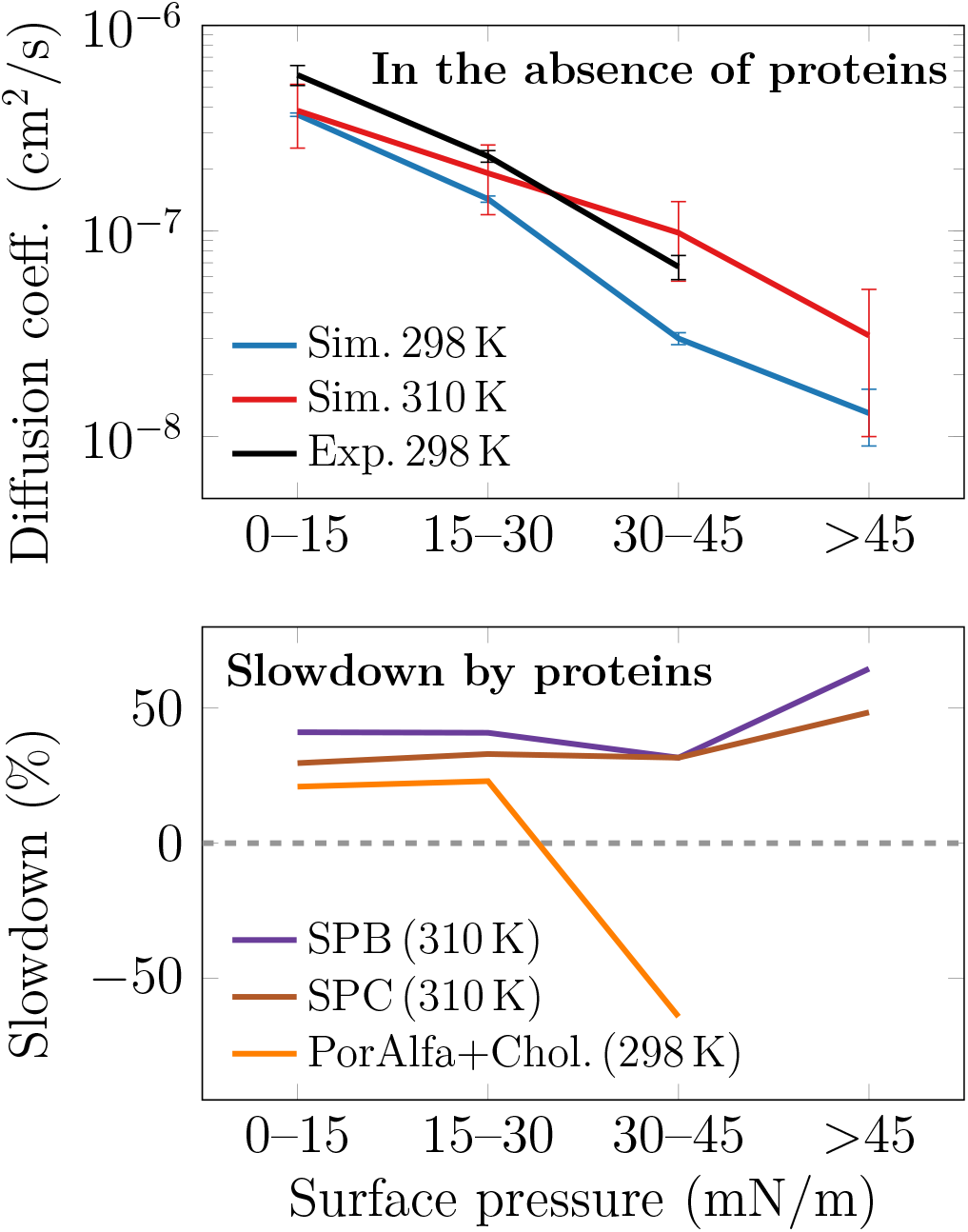
Diffusion coefficients of lipids as a function of surface pressure. Top: diffusion coefficients in protein-free monolayers from both simulations (“Sim.”) and experiments (“Exp.”). Bottom: effect of surfactant proteins on diffusion given as percentages of slowdown (positive values) or speedup (negative values). The data from simulation are averaged over all lipids, whereas those from the experiment were measured using *z*-scan FCS with fluorescent DOPE-Atto633 dye. Here, “PorAlfa+Chol.” stands for poractant alfa with 10% of CHOL added.

The increase in surface pressure leads to a significant slowdown of diffusion. In simulations, we averaged over the motion of all lipid species, whereas the *z*-scan FCS measurements probe the motion of Atto633-labeled DOPE lipids that are present at a low concentration. Thus, both methods should be comparable as they measure single-lipid diffusion coefficients. As demonstrated in the top panel of Fig. 3, the agreement between the values for protein free NoP systems extracted from simulations and experiments is excellent both in terms of the diffusion coefficient values, as well as their decreasing trend due to compression. The slowdown from low (0–15 mN/m) to high (30–45 mN/m) pressure is almost an order of magnitude. Further compression to pressures above 45 mN/m in simulations leads to a decrease by almost another order of magnitude, yet this region cannot be probed using experiment due to monolayer collapse. The simulations suggest that the diffusion coefficients and their trends at 298 K and 310 K are fairly similar.

As shown in the bottom panel of Fig. 3, the effect of proteins at low and intermediate surface pressures is to decrease lipid diffusion coefficients, and this behavior is reproduced by both experiments and simulations. In simulations, the slowdown is 30–40%, and slightly higher for SPB than for SPC systems, whereas experiments suggest a slowdown of 20–25%. Such effects are observed also for lipid bilayers at low protein concentrations. ^77^ However, at a higher pressure of 30–45 mN/m, things change: Diffusion in simulations slows down by 50% or more upon the insertion of SP-B or SP-C. Strikingly, diffusion in experiments is faster in the protein-containing poractant alfa than in the protein-free quaternary mixture. One possible explanation for this discrepancy is that we only have one protein per monolayer in our simulations, whereas the poractant alfa used in experiments contains 1% of SPs, which could thus collectively perturb the monolayer packing and increase lipid mobility by, *e.g*., the formation of SP-B oligomers. ^15,24^ Moreover, our synthetic protein-free mixture is much less complex than poractant alfa in terms of lipid composition. Still, our model reproduces substantially well both the lipid diffusion coefficients, their decrease due to compression, and the effect of proteins at low pressures, *i.e*. in the uniform L_e_ phase.

In conclusion, our simulation model reproduces the experimental behavior of the multicomponent monolayer under non-equilibrium conditions and avoids common methodological pitfalls. ^65,66^ Moreover, the effects of SPs are qualitatively, and to a large extent also quantitatively, reproduced by our simulation model.

### Hydrophobic Surfactant Proteins Reside at Different Depths in the L_e_ Phase

MD simulations have the ability to resolve the locations of proteins in the surfactant monolayers, thus providing a molecular level explanation for the trends observed in the isotherms in Fig. 2. Earlier experimental and simulation studies have focused on the location and orientation of SP-B and SP-C in model bilayers, ^40,40–43,69,70^ yet with somewhat contradicting results. However, no studies have reported the vertical positioning and orientation of SPs in PSurf monolayers. Our MD simulations readily provide this information as a function of surface pressure.

We first extracted the orientation of SP-C in the PSurf monolayer as a function of monolayer compression. In the SPC systems, the SP-C protein was found to remain almost parallel to the monolayer surface at an average angle of 95.3±4.4° relative to the monolayer normal (Fig. S5). This agrees with the SP-C orientation in bilayers resolved recently by scattering experiments, ^40^ and contradicts with the earlier data suggesting a transmembrane orientation. ^41,42^ Curiously, CHOL has been demonstrated to affect SP-C orientation in bilayers, ^43^ suggesting that it might depend on membrane order. However, it seems that the thickness of the PSurf monolayer or the surface pressure applied to the interface has little to no effect on the tilt angle of SP-C. The equilibrium tilt angle of SP-C was further validated by an additional set of simulations at constant APL in which SP-C was initially placed at different angles parallel and perpendicular to the monolayer normal at APL of 90 Å^2^. SP-C achieved the equilibrium tilt angle parallel to the monolayer plane within the first 10 nanoseconds, and remained at the same angle through the 100–500 ns simulations (Fig. S5).

Moving on, we extracted the density profiles of SP-B and SP-C in the simulations as a function of surface pressure (see Methods in the SI for details). These data are plotted in Fig. 4 as a 2D map so that a vertical slice at any surface pressure value would provide the typical density profile along the monolayer normal at that surface pressure. The profiles are also aligned so that phosphorus atoms remain at the same depth, rendering it straightforward to evaluate the positioning of SPs with respect to the air–water interface. The curves show the positions between which each molecule type displays density that is larger than 5% of its maximum density.

**Figure 4:**
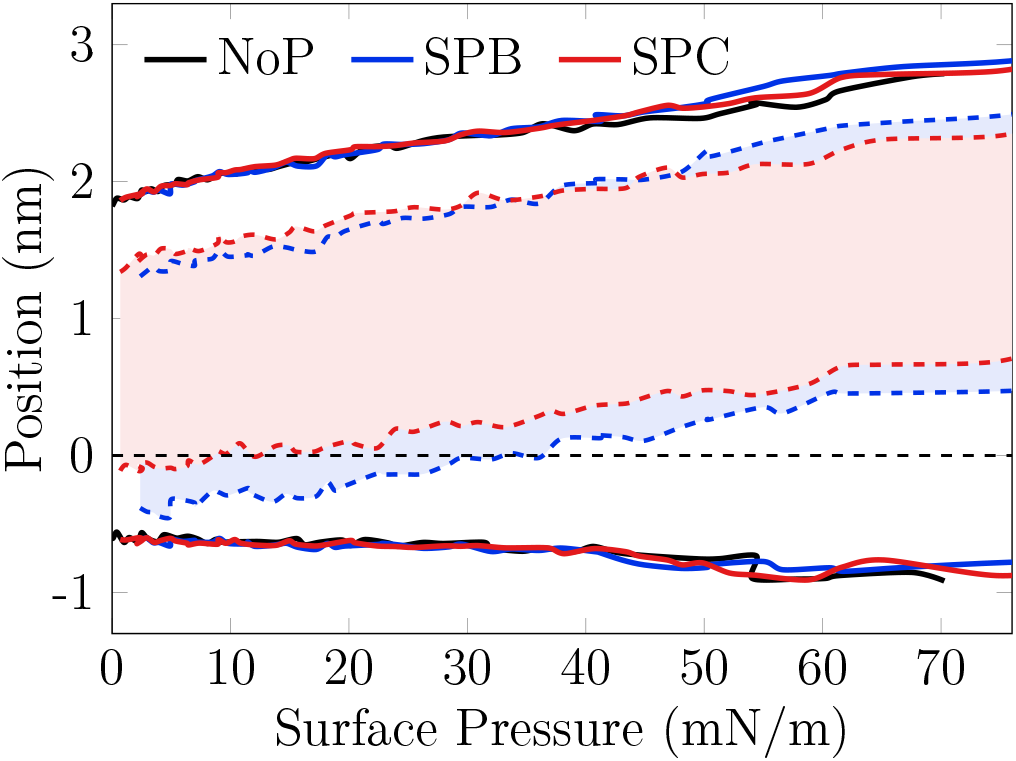
The average thickness of the monolayers and the position of the surfactant proteins as a function of surface pressure. The curves show the positions (normal to the monolayer) at which the respective density reaches 5% of its maximum value. The solid lines, representing the phospholipids in the different systems, are essentially identical indicating that the proteins do not have a significant effect on the monolayer thickness. The shaded area between the dashed blue and red lines show the position of SP-B and SP-C, respectively. The zero position is defined as the position of the phosphorous atoms of the phospholipids.

Both the protein-free systems (“NoP” in Fig. 4) and the systems with proteins (“SPB” & “SPC”) demonstrate that the monolayer gets significantly thicker upon compression, from ~2.5 nm to ~4 nm. This is coupled to the decrease of the average tilting of the lipid acyl chains (Fig. S7), although a fraction of the lipids might show a characteristic tilt when part of the L_c_-like clusters. ^11^ Most of this thickening of more than 1 nm occur in the acyl chains region, corresponding to an increased chain ordering. The head groups also contribute a little by extending towards the aqueous phase. Considering the individual lipid components in the model surfactant, across all surface pressures, the POPG head group seems to reside slightly below those of DPPC and POPC (Fig. S6). Still, on average all the phospholipids show similar trends, whereas CHOL shifts slightly away from the interface upon compression and resides in the acyl chain region at higher surface pressures.

The presence of proteins (“SPB” & “SPC” in Figs. 4 and S6) has a lesser effect on the monolayer thickness or the relative positioning of the different lipid types. However, compression has a significant effect on SP positioning in the PSurf monolayers. SP-B initially resides at the interface and partially above the phosphate groups in the low surface pressure L_e_ phase, resembling the positioning resolved experimentally in bilayers. ^40^ However, SP-B re-positions itself into the lipid acyl chain region well below the phosphate groups upon compression. SP-C remains below the phosphate level even in the L_e_ phase—again similar to its behavior in the bilayer^40^—yet obtains a similar positioning as SP-B in the acyl chain region upon compression. These findings suggest that the ability of SP-B to increase surface pressure results from its presence at the interface in the L_e_ phase. SP-C is less present in this region and therefore affects lipid head group packing and consecutively surface pressure to a smaller extent. However, this partitioning alone does not explain the major pressure increase caused by SP-B in the L_c_ phase (Fig. 2). In this regime, a viable explanation is that SP-B causes significant perturbation in the packing of the L_c_ phase. Indeed, as demonstrated in Fig. S8, both SPs greatly affect the tilt of lipid acyl chains in their vicinity with SP-B inducing a greater effect. Moreover, POPG demonstrates the highest tilt, indicating that it might be the most affected by the SPs (Fig. S7).

The perturbation of the monolayer structure also relates to the phase-preference of the SPs. We clustered selected atoms in the lipid acyl chains and CHOLs using the DBSCAN algorithm (see Methods in the SI for details), and the found tightly-packed clusters were associated with the L_c_ phase. We then calculated the fraction of lipid chains in the L_c_ phase as a function of the distance from the protein. The calculation was performed on four distinct surface pressure regimes, and the resulting distributions are shown in Fig. 5.

**Figure 5:**
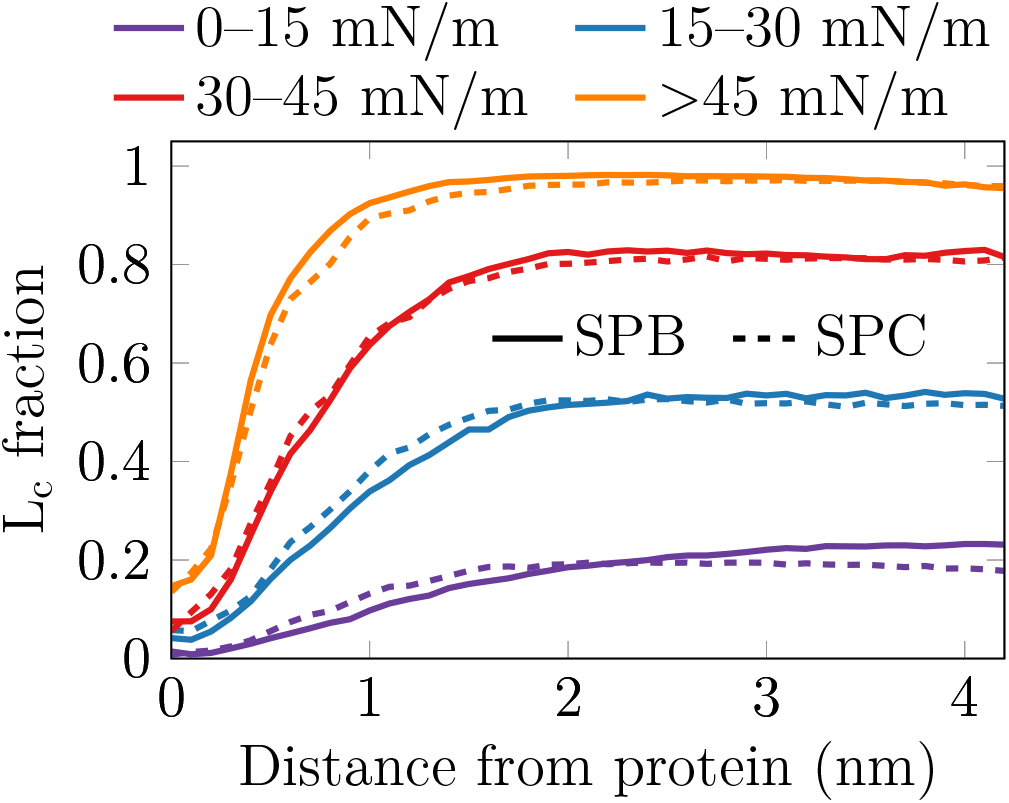
The fraction of the lipid acyl chains that show L_c_-like packing as a function of the shortest distance from the protein. Data are shown for 4 different surface pressure regimes. The L_c_-like packing is detected using DBSCAN algorithm performed on the 10_th_ carbons in the lipid acyl chains, as well as the C14 atom of CHOL that resides on the same depth.^11^

It is evident that both SPs prefer to reside in the L_e_ phase at all surface pressure regimes. Actually, the L_c_ fraction near the protein is always lower than far from it, indicating that the SPs actually induce the L_e_ phase in their immediate vicinity. Depending on the surface pressure, the effect seems to range between ~1 nm in the high-pressure regime and ~2 nm in the low-pressure regime from the protein surface. This suggests that in a monolayer with a realistic concentration of SPs (10 wt-% of total protein^4^), the SPs could greatly perturb the overall structure of the monolayer and thus increase the surface pressure. Moreover, the ability of SPs to partition the L_c_ phase into smaller islands^28,30–34^ is likely caused by this perturbation effect. Although its range is very similar for both proteins, the overall perturbation by SP-B will be greater due to its larger size, in line with the effects of SPs on surface pressure (Fig. 2).

To verify the observed effects of SPs on the lateral organization of the PSurf monolayer, we performed atomic force microscopy (AFM) measurements of films transferred to a mica surface. For these measurements, we used the same lipid mixture as in our simulations, and either SP-B or SP-C was added (see Methods for details). The corresponding AFM data for the protein-free systems were reported in our previous work. ^11^ The AFM images are assembled in Fig. 6.

**Figure 6:**
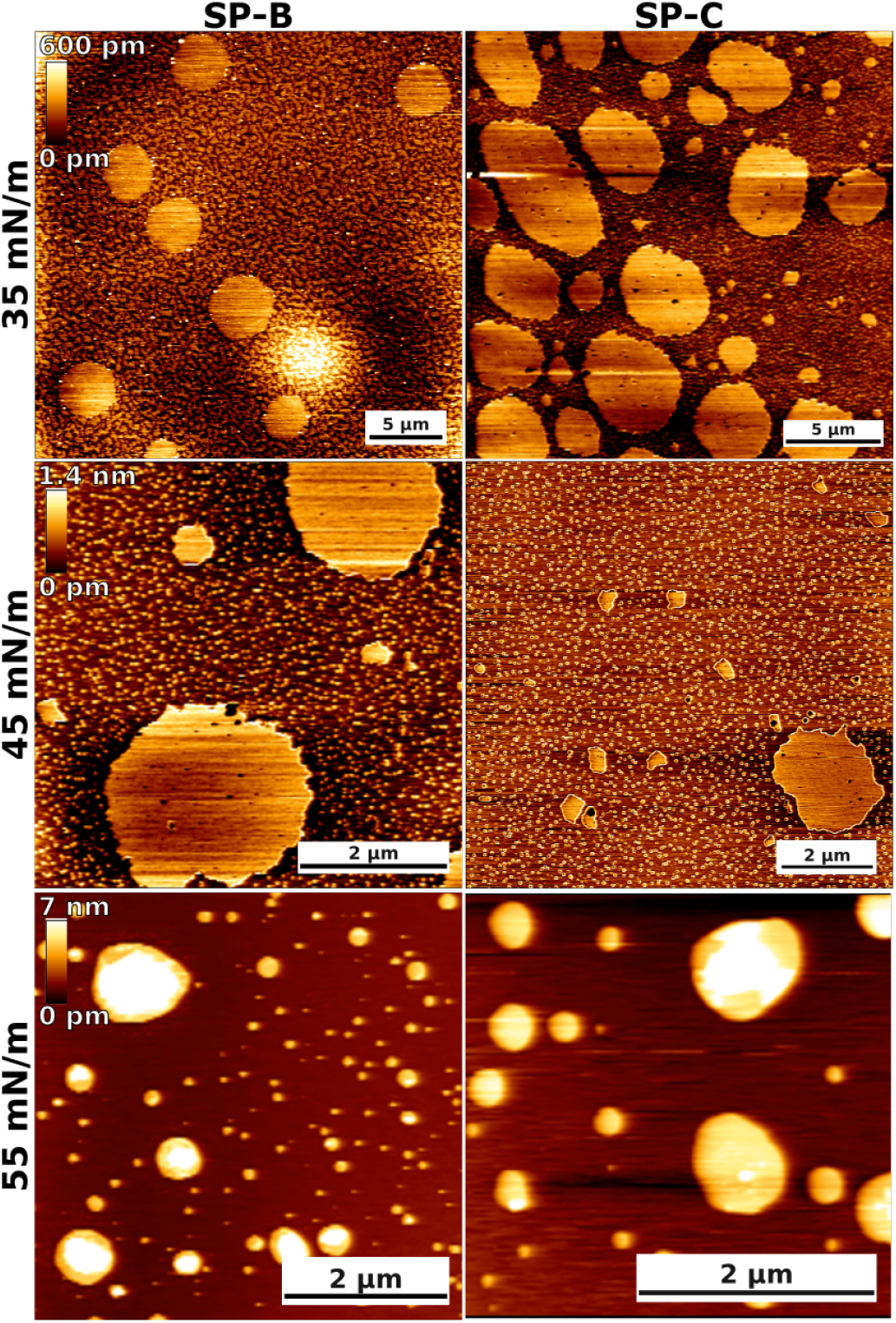
Atomic force microscopy imaging of model pulmonary surfactant monolayers with SP-B and SP-C at different surface pressures. The scale bars equal to 5, 2, and 2 μm at 35, 45, and 55 mN/m, respectively.

At a surface pressure of 35 mN/m, the monolayers contain large and roundish domains that are ~0.2–0.5 nm thicker than the surrounding regions, so we assign them to the L_c_ phase. In addition, the monolayers contain smaller-scale heterogeneity with more elongated and irregularly shaped L_c_ domains which have not coalesced into large domains. The shape and small size indicate that the line tension associated with their boundaries is not significant. While the qualitative behavior of the systems with SP-B and SP-C are similar, it seems that the thickness difference between the L_e_ and L_c_ regions is smaller in the presence of SP-B (~0.2–0.3 nm) than in the presence of SP-C (~0.4–0.5 nm). Moreover, the area coverage of the L_c_ phase seems to be somewhat larger in the case of SP-B, whereas the boundaries of the L_c_ phase are sharper in the presence of SP-C. Together, these findings suggest that SP-B possibly partitions more to the L_c_ phase and renders the two phases less distinct by inducing disorder within the L_c_ phase. On the other hand, SP-C is possibly excluded from the L_c_ phase altogether, allowing its lipid chains to fully extend for maximal thickness difference. The molecular level explanation for this difference is not evident from our simulation data.

The same trends are largely present at a surface pressure of 45 mN/m. In our previous work, we observed the L_c_ phase to cover most of the protein-free monolayer with only small L_e_-like islands present at this pressure. However, in the case of SP-B or SP-C, the quasicontinuous L_c_ phase is split into smaller L_c_ islands within a percolating L_e_ phase. This behavior is in line with the ability of the SP-B and SP-C to break down the L_c_ phase observed in our simulations and earlier experiments.^28,30–34^ Moreover, the looser packing of the L_e_ phase means that this breakdown leads to an increase in surface pressure, which is visible in the surface pressure–area isotherms from simulations (Fig. 2) and experiments.^33–36^ The larger effect of SP-B on the surface pressure could possibly results from its larger partitioning to the L_c_ phase and the perturbation of the packing therein.

At a surface pressure of 55 mN/m, the monolayer has collapsed for both proteins with protrusions of multiple nanometers. Similar behavior was also observed in the protein-free case.^11^

### Surfactant Proteins Engage in Specific Lipid–Protein Interactions

In addition to the physical properties of the PSurf monolayer, MD simulations can also resolve the specific lipid–protein interactions. These interactions are challenging targets for experimental approaches due to their relatively weak and transient nature. Moreover, the labels required by fluorescence approaches would likely substantially perturb these interactions. We first extracted the contacts between non-hydrogen atoms of the lipids with the proteins, and normalized these values based on the number of possible contact partners. We then calculated the average values of these contacts as a function of surface pressure. These data are shown in Fig. 7A.

**Figure 7:**
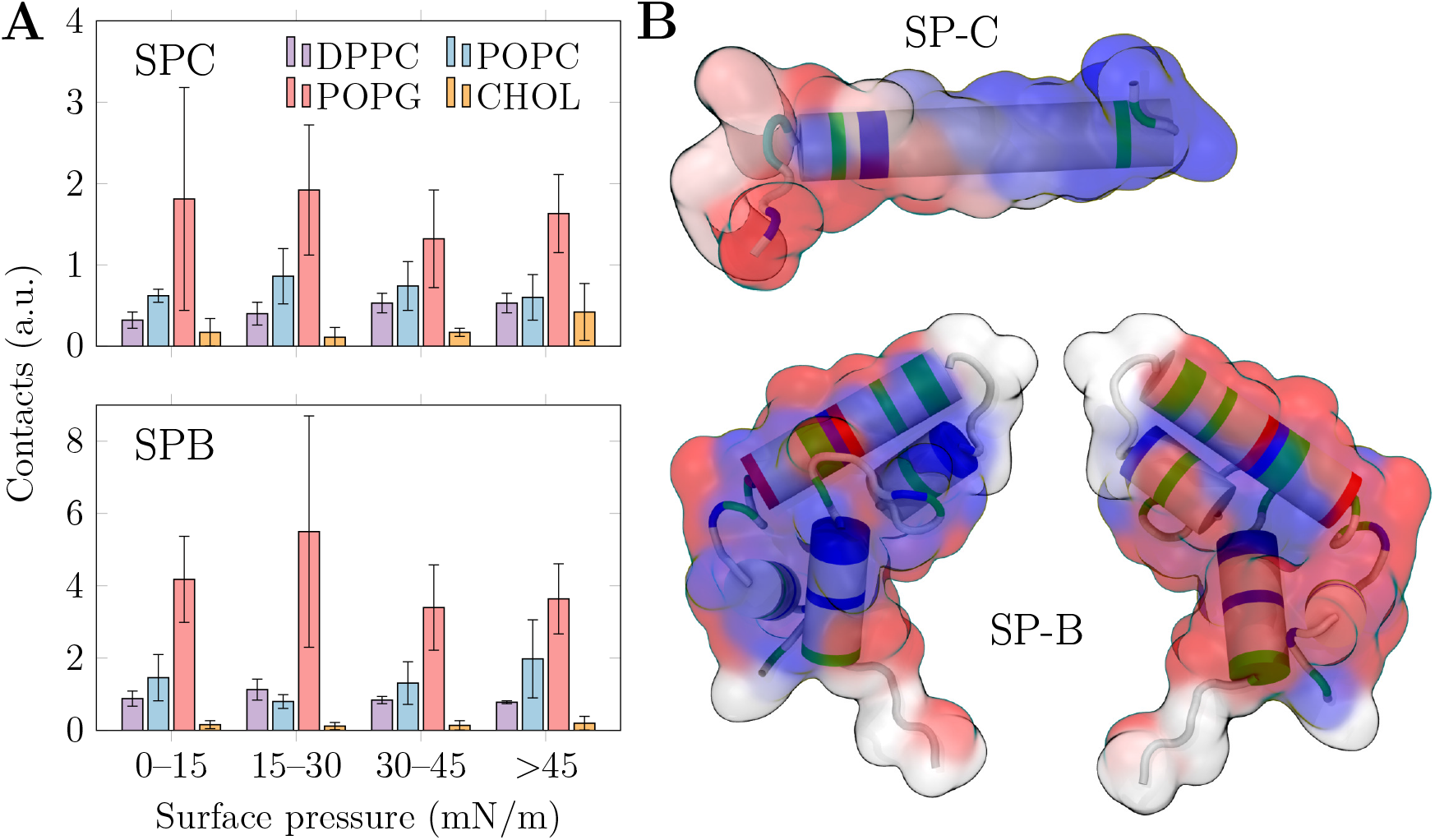
The lipid–protein interactions between SPs and monolayer lipids in MD simulations. A) Effect of surface pressure on preferential protein–lipid contacts (cutoff 0.3 nm). The data are shown for four ranges of surface pressure. The error bars show standard error, extracted from the mean values of four individual monolayers (2 per simulation system, 2 repeats). B) Interactions with POPG at low (0–15 mN/m) surface pressure mapped to the protein structures. On the surface representation blue, white, and red indicate low, medium, and high interaction frequency with PG, respectively, and on the cartoon representation the types of the amino acids are shown as blue, red, green, and white for basic, acidic, polar, and other, respectively. The two sides of SP-B are shown separately. Note that the palmitoyl chains attached to SP-C residues 5 and 6 are omitted from the visualization.

The trends in Fig. 7A demonstrate that POPG is a preferred interaction partner for both SPs and at all surface pressure ranges in the simulations. For SP-B, the preference for POPG is 4-fold compared to that of POPC or DPPC, whereas for SP-B the ratio is only 2 or even smaller. The surface pressure has little effect on the trends in general, which is likely due to the slow lateral diffusion of lipids at higher surface pressures. Therefore, the preferential interactions adopted at low surface pressures likely remain dominant upon compression to higher pressures.

The observed tendency of SP-B to preferentially interact with PG is well-established,^15,28,50^ although there have been some contrasting reports as well.^38,51^ For SP-C, the situation is less clear, and experiments typically have not observed a major preference for POPG. ^28^ This goes hand in hand with the relatively small preference of SP-C to interact with POPG in our simulations. Curiously, CHOL has little role in interactions even with SP-C. This agrees with earlier reports, ^27,28^ and signals that the mechanism through which CHOL affects SP-C orientation in bilayers^43^—if real—is likely a membrane-mediated effect caused by the overall thickening and ordering of bilayers by CHOL^79^ instead of specific lipid–protein interactions. The lack of SP–CHOL interactions is also evident in the snapshots shown in Figs. 1 and S1.

Both SP-B and SP-C proteins slightly prefer interactions with POPC over DPPC. All in all, these data agree with SP-B and SP-C remaining in the L_e_ phase even when L_c_-like clusters form in the monolayer. Although there are no studies in which the partitioning preference of SPs was studied in monolayers, the results go hand in hand with data obtained from lipid membranes in which the SPs prefer the more disordered L_d_ phase over the ordered L_o_ one.^17^

Having established the general lipid preferences of the SPs, we looked into the specific lipid–protein interactions by calculating the mean number of contacts with different lipid types for each residue in the SPs. We performed the analyses on the different surface pressure regimes, and normalized the data based on the composition of our model PSurf. The complete data are visualized in Figs. S9 and S10 in the SI for the SPC and SPB systems, respectively. Following Fig. 7A, no residues show specific interactions with CHOL nor DPPC for either protein. Out of the two PCs, SP-C demonstrates an overall preference to interact with POPC in all surface pressure regimes, but this cannot be pinpointed to any particular residues. For SP-B, the interactions with POPC manifest themselves only in the compressed monolayer, suggesting that they are also related to the the partitioning preference of SP-B rather than to any specific interactions.

As indicated by Figs. S9 and S10, the interactions between SPs and POPG are more specific. For SP-C, these interactions are concentrated on the first 15 residues in the N-terminal where the palmitoylated cysteine residues are also located. The highest number of contacts with POPG is observed for Asn9. These interactions are best highlighted at low surface pressure, where lipids can freely diffuse to form these interactions. However, at higher surface pressures these preferences are somewhat lost. In the low surface pressure regime, SP-B demonstrates preferential interactions with POPG at Arg17, which also promotes contacts with the nearby Leu10 and Leu14. Towards the C-terminal, Arg76 and Arg72 and the nearby Val74 show increased contacts with POPG, as do Arg64 and the nearby Met65 and Leu61. At larger pressures, the contact preferencies vary somewhat, yet they most important POPG contact partners are always concentrated around the few arginine residues listed above.

To better visualize the location of the POPG interaction sites on the SPs, we mapped the low-pressure data in Figs. S9 and S10 to the protein structure in Fig. 7B. The surfaces show the tendency to interact with POPG (blue:low, white:medium, red:high), whereas the cartoon representation shows the types of the amino acids (blue:basic, red:acidic, green:polar, and white:other). For SP-C, the interactions are concentrated in the N-terminus with two arginine residues. For SP-B, not all acidic residues are involved in the interactions. Instead, the larger size of the protein limits the contacts of residues in the protein core from interacting with any lipids, and the POPG interactions are concentrated on the protein edges. Moreover, the two sides of SP-B show different interactions due to the preferred orientation of SP-B on the monolayer. ^15^

## Conclusions

We have used multi-microsecond atomistic non-equilibrium MD simulations to study the effect of hydrophobic surfactant proteins on pulmonary surfactant monolayers, the positioning and phase-preference of these proteins in the surfactant, and their specific interactions with lipids. All these analyses were extracted across a range of compression states (*i.e* surface pressures), corresponding to the entire breathing cycle. We used the recently suggested model combination^11,65,66^ to accurately capture the interfacial physics. Moreover, we performed *z*-scan FCS experimental measurements to validate that our simulation approach correctly captures the effects of compression and proteins on lipid mobility, as well as AFM imaging that confirms the role of proteins in promoting the L_e_ phase.

Our simulations reproduced the experimentally observed effects of surfactant proteins on the isotherms and diffusion coefficients. The pressure-increasing ability of SPs^34,36^ was linked to their penetration to the head group region in the L_e_ phase. During compression, SP-B re-positioned itself to below the head group region. Therefore, the local perturbation of acyl chains and the induction of the L_e_ phase in its vicinity explain the increased surface pressure in the compressed L_c_-like monolayer. The smaller increase in the surface pressure due to SP-C could result from its preferential positioning below the head group region and parallel to the monolayer. In addition, AFM imaging suggests that the difference in the thickness of the L_c_ and L_e_ phases decreased with SP-B, indicating that a fraction of SP-B could partition to the L_c_ phase and decrease its packing density and thus increase the surface pressure of the monolayer.

Both SPs showed interaction preference only towards POPG in our 4-component lipid mixture, yet this tendency was substantially larger for SP-B, in line with experimental evidence. ^28,50^ Not surprisingly, arginine residues preferentially interacted with POPG. In SP-C, these residues are located in the N-terminus, whereas in SP-B the geometry of the protein also affects the interactions, which are concentrated around arginine residues on the sides of the protein.

All in all, our study represents a detailed look into protein–lipid interactions in our *in silico* model for the pulmonary surfactant. Understanding the roles of its lipid and protein components is currently a bottleneck in the development of improved synthetic surfactants for the treatment of newborn respiratory distress syndrome *via* surfactant replacement therapy.^80^ Moreover, the surfactant system poses both a health hazard and therapeutic potential, as it provides an efficient route to the body for not only undesired pollutants and pathogens,^81^ but also for surfactant-coated^82^ and nebulized drugs,^83^ rendering its role for human health and well-being significant. Computer simulations have the potential to contribute to these challenges, and we believe that the present manuscript presents a key step on the path towards more accurate and realistic PSurf models.

## Methods

### Atomistic Molecular Dynamics Simulations

We extended our previous work on monolayers with the same lipid mixture,^11^ yet this time we included SP-B and SP-C proteins in the simulations and opted for dynamic simulation conditions. The SP-B model was obtained from our previous work,^15,24^ whereas the SP-C model was generated as an alpha-helical peptide with two palmitoyl chains inserted in CHARMM-GUI.^84^ A monolayer simulated for 1000 ns with an APL of 90 Å^2^ was obtained from our previous work. ^11^ The SPs were incorporated in the monolayers by placing the proteins within 1 nm of the lipids either on the side of the headgroups or the acyl chains and slowly letting them integrate into the monolayers. Simulations initiated from these two conformations served as two independent replicas, both with two monolayers and thus two SPs.

The CHARMM36 topologies^72,73^ were downloaded in GROMACS formats,^85^ and the protein-containing systems were subjected to the standard equilibration protocol. ^85^ After equilibration, the monolayers with the SPs were simulated without restraints for an additional 500 ns each. For the SP-free system, we used our existing simulation setup. ^11^ The compression/expansion simulations were performed using the deform keyword in GROMACS with a linearly changing edge of a square monolayer area so that the area per lipid changed from 45 Å^2^ to 90 Å^2^ (or *vice versa*) during the simulation time of 5 μs. For NoP, 5 μs compression and expansion simulations were performed, whereas for SPB and SPC only 5 μs compression simulations were performed. The compression/expansion rate of the simulation box was linear in time, resulting in a quadratic scaling of the area of the square monolayer. All systems were simulated at the physiological temperature of 310 K, yet the NoP system was also simulated at 298 K (both compression and expansion) to evaluate the effect of temperature on phase behavior and hysteresis.

Following experimental approaches, the protein area in SPB and SPC systems was not excluded in any way in the definition of APL, since the monolayer compression state heavily affects the transverse protein location and thus renders any such attempts ambiguous. Still, APL is only considered in our surface pressure–area isotherms, whereas other results are reported as a function of surface pressure instead. In the analyses of protein-containing systems, we averaged the results over two replicas both with two monolayers and thus two SPs for a total of four independent samples. Standard error was obtained as the standard deviation of the mean values extracted for these independent samples, and used as the error estimate. Apart from the dynamic box size, the simulation parameters and force fields followed our earlier work^11^ and thus the standard parameters for CHARMM force fields in GROMACS.^85^ The simulation parameter file (.mdp) can be downloaded together with all simulation outputs from the Zenodo repository at DOI:DOI:10.5281/zenodo.6817824). The analyses were performed using the tools included in the GROMACS^86,87^ simulation package as well as in-house tools.

### Materials for Z-scan Fluorescence Correlation Spectroscopy Measurements

1,2- dipalmitoyl-*sn*-glycero-3-phosphocholine (DPPC), 1-palmitoyl-2-oleyl-*sn*-glycero-3-phosphocholine (POPC), 1-palmitoyl-2-oleoyl-*sn*-glycero-3-phospho-(1’-rac-glycerol) (POPG), and CHOL (ovine wool) were purchased from Avanti Polar Lipids (Alabaster, AL). An extract of natural porcine lung surfactant, poractant alfa (Curosurf, Chiesi Farmaceutici, Parma, Italy), enriched with CHOL (10 wt%) was used in control experiments. 1,2-dioleoyl-sn-glycero-3-phosphoethanolamine labeled with Atto633 (DOPE–Atto633) was obtained from ATTOTEC (Siegen, Germany). Organic solvents of spectroscopic grade used for the preparation of lipid working solutions were supplied by Merck (Darmstadt, Germany). Phosphate-buffered saline (PBS) (Sigma-Aldrich, St. Louis, MO) prepared with Mili-Q water (Millipore, USA), with addition of 0.2 mM ethylenedinitrilotetraacetic acid (EDTA) (Sigma-Aldrich, St. Louis, MO), was used as a subphase in the experiments. All chemicals were used without further purification.

### *Z*-scan Fluorescence Correlation Spectroscopy

The *z*-scan FCS measurements were performed on the “NoP” mixture (60/20/10/10 mol% of DPPC/POPC/POPG/CHOL) and on the extract of natural porcine lung surfactant supplemented with 10 wt% of CHOL that is filtered out during the production of poractant alfa. Langmuir μtrough XS setup (Kibron, Helsinki, Finland) was placed on an inverted confocal fluorescence microscope (Olympus, Hamburg, Germany) equipped with the water-immersion UPlanSApo 60× objective (NA 1.2, WD 0.28 mm, Olympus). Lipid mixture in chloroform was spread over a subphase filling stainless steel trough with PTFE edges and an in-house modified glass window with a Hamilton microsyringe. The fluorescent probe, DOPE-Atto633, present in a molar ratio of 1:300,000 to other lipids, was excited with the pulsed laser (635 nm; LDH-D-C-635, Pico-Quant). Fluorescence signal, after passing trough Z473/635 dichroic (Chroma, Rockingham, VT), 100 nm pinhole and 685/50 band-pass emission filter (Chroma, Rockingham, VT), was collected with a single photon avalanche diode (SPAD). After chloroform was allowed to evaporate from the interface (~10 min), the compression of spreaded film was initiated with the symmetrical movement of two Delrin barriers controlled by FilmWare software at a constant rate of 3.92 Å^2^/molecule/min. During surface pressure–molecular area (Π–APL) isotherm recording with an ultra-sensitive surface pressure sensor with the DyneProbe, compression was stopped at selected surface pressures within the range from 5 to 45 mN/m with a step of 5 mN/m for 5 min after which a *z*-scan FCS measurement was performed. The monolayer was scanned verticaly every 200 nm in up to 20 steps. More detailed description of *z*-scan FCS method and data analysis can be found elsewhere. ^88^ For each system, the experiment was done in triplicate at 298 K. Subphase temperature was controlled with a temperature plate placed under the trough and its evaporation during the measurement was compensated with constant controlled addition of subphase with a peristaltic pump from the outside of barriers.

### Langmuir–Blodgett Transferred Monolayers and Atomic Force Microscopy

We employed a specially designed Langmuir–Wilhelmy trough (NIMA Technology, UK). Compression isotherm assays were performed and surface pressures were kept constant at constant temperature, as described previously by Dohm et al..^89^ The lipid mixture used was the same as for the simulations and FCS studies dissolved in chloroform:methanol 2:1 (v:v). The solution contained 1% of SP-B or SP-C. Surfactant proteins were purchased from Seven Hills Bioreagents (Cincinnati, Ohio). The Langmuir trough has ribbons instead of Teflon barriers and allows for a maximum area of 312 cm^2^ and a minimum of 54 cm^2^. The continuous Teflon-coated ribbon closes its area where the lipids are deposited by moving symmetrically two barriers, each holding 2 Teflon barrels. Pressure recordings are done by an electronic pressure sensor, where a piece of cellulose is hanging from a copper hook. The measurements are done employing the Wilhelmy technique with an estimated error of ±1 mN/m among different isotherms. A minimum of triplicates at 298 K were done. Lipid monolayers were transferred onto freshly cleaved muscovite mica substrate (Plano GmbH, Wetzlar, Germany) as described by Brown et al.. ^90^ The transfer ratio was 1; no compression or expansion of the monolayer took place during the transfer. In order to achieve an equilibrated monolayer at the air–liquid interface, the lipid sample was deposited carefully and let spread until a minimum surface pressure of ~0–1 mN/m was observed. After 10 min of monolayer equilibration, the film was compressed until the desired surface pressure was reached at a compression speed of 50 cm^2^/min. Before the transfer started, the film was again equilibrated for 5 min at constant pressure, and monolayers were deposited onto previously submerged mica. The lifting device was raised in the vertical plane out of the buffered aqueous subphase at a speed of 10 mm/min at constant pressure. Three to 5 independent experiments were carried out. Langmuir–Blodgett supported monolayers’ topographical images were taken using an atomic force microscope (JPK NanoWizard, JPK Instruments, Berlin, Germany), employing in both cases Silicon-SPM cantilevers (Nanosensors, NanoWorld AG, Neuchatel, Switzerland). The AC mode in air was selected for monolayers. The scan rate was ~1 Hz for all AFM images. At least three different supported monolayer systems were assessed, and each sample was imaged on a minimum of three different positions. Image processing of AFM data was done using the JPK imaging software package provided by JPK Instruments.

## Acknowledgement

We thanks CSC–IT Center for Science for computational resources. J.L. thanks the Sigrid Jusélius Foundation for funding. A.O. and L.C. were supported by Czech Science Foundation (grant no. 21-19854S). I.V. thanks the Helsinki Institute of Life Science (HiLIFE) Fellow program, Human Frontier Science Program (HFSP, project no. RGP0059/2019), Sigrid Jusélius Foundation, and Academy of Finland (project ID: 335527, 331349) for financial support. J.B.S acknowledges support from Bill & Melinda Gates Foundation and BBSRC (INV-016631 and BB/V019791/1, respectively); also from the Commonwealth and the Integrated Biological Imaging Network (IBIN), a Technology Touching life MRC Network. M.J. thanks the Academy of Finland (Postdoctoral grant no. 338160) and the Emil Aaltonen foundation for funding.

## Supporting Information Available

Methodological details and additional results.

