## Supporting Information for "Surfactant Proteins SP-B and SP-C in Pulmonary Surfactant Monolayers: Physical Properties Controlled by Specific Protein–Lipid Interactions"

### Analysis Methods

**SP-C Tilt Angle** The tilt angle of SP-C was calculated as the the angle between the $z$ axis (normal to the monolayer plane) and a vector joining residues Lys11 and Leu32 $C_\alpha$ atoms using the `gmx gangle` tool provided with GROMACS.[1] The tilt angles were averaged over the four repeats and over the length of the simulations, and the error estimate is given as the standard error.

**Lipid Acyl Chain Tilt Angle** The phospholipid acyl chain tilt angle was calculated as the angle between the $z$ axis (normal to the monolayer plane) and a vector joining the 1st and 16th carbon (C21 and C216) in the *sn*-1 fatty acid chain using the GROMACS tool `gmx gangle`. The lipid acyl chain tilt angles were averaged per lipid type over the four repeats for the SPB and SPC systems and over the two repeats for the NoP systems. The distance of a lipid from the protein was calculated with the GROMACS tool `gmx mindist` every 1 ns as the minimum distance of a lipid phosphorus atom from the protein.

**Surface Pressure–Area Isotherm** Monolayer surface pressure $\Pi$ as a function of APL $A$ was calculated as $\Pi(A) = \gamma_0 - \gamma(A)$, where $\gamma$ and $\gamma_0$ are the surface tensions of the surfactant-covered and surfactant-free air–water interfaces, respectively. The surface tension $\gamma$ was calculated as $\gamma = L_z \times (P_\mathrm{N} - P_\mathrm{L})/2$, where $L_z$ is the length of the simulation box $z$ axis normal to the interface, $P_\mathrm{N} = P_\mathrm{zz}$ and $P_\mathrm{L} = (P_{xx} + P_{yy})/2$ are the normal and planar components of the pressure, respectively, and the factor $1/2$ accounts for the two interfaces in each simulation system. The values were obtained with the GROMACS tool `gmx energy`. The area of the monolayers was extracted from the size of the simulation box as $L_x \times L_y$, where $L_x$ and $L_y$ are the lengths of the simulation box $x$ and $y$ axes, respectively. The trajectories were analyzed in 100 ns sections and the values were calculated as an average over the section. The results were then averaged over the repetitions and the error estimate was calculated as standard error.

**Lipid Diffusion Coefficients** The lateral diffusion of lipids was quantified using displacement distributions. To this end, we extracted the displacements of lipid centers of mass over 10 ns intervals. The values extracted from different surface pressure regimes (0–15, 15–30, 30–45, and >45 mN/m) were histogrammed. For simplicity, displacements across periodic boundaries were discarded, yet this has no effect on the shape of the distribution. The distributions were normalized and fitted by[2]

$$P(r, \Delta) = \frac{r}{2D\Delta} \exp\left(\frac{-r^2}{4D\Delta}\right), \tag{1}$$

where $r$ is the length of the displacement, $\Delta$ the time interval, and $D$ the diffusion coefficient. We chose $\Delta$=10 ns to avoid probing anomalous diffusion at short timescales,[3] while also probing the desired surface pressure interval; Using larger $\Delta$ value would mean that the displacements start and end at very different surface pressures in the dynamic simulation. The diffusion coefficients were measured for two (NoP system, two monolayers) or four (SPB and SPC systems, two replicas with two monolayers) monolayers and averaged. Standard error of these independent samples was used as the error estimate.

**Time-Resolved Density Profiles** The thickness of the monolayer and the insertion depth of the proteins into the pulmonary surfactant were determined by calculating the density profiles of the components along the monolayer normal ($z$ axis) with the GROMACS tool `gmx density`. The simulation box was divided into 1 Å slices and the densities were calculated every 10 ns. These profiles were aligned at the peak density of the phosphorous atoms of the phospholipids and the results were averaged over all monolayers. The averaged lipid and protein densities were set to begin and end at $z$ values where the density crossed 5% of its maximum value.

**$L_e$ Phase Fraction** To classify the lipids into $L_c$ or $L_e$ phases, we applied the DBSCAN algorithm[4] to the coordinates lipid acyl chains and cholesterols. For lipids, we considered the $10^{th}$ carbons in the lipid acyl chains, whereas for cholesterol the "C14" atom—residing at a similar level as the chosen acyl chain carbons—was chosen. These coordinates were projected to the plane of the monolayer, and the DBSCAN was performed with a cutoff of 0.71 nm and a number of neighbours set at 6, following our earlier work.[5] These parameters lead to the expected result of almost 100% of $L_c$ coverage far from the protein in the most compressed systems.

For each acyl chain or cholesterol included in the analysis, its closest distance to a protein alpha carbon was extracted, and the types of the acyl chains ($L_c$ or $L_e$) were binned to a histogram based on this distance. After averaging over the two simulations and the two monolayer in both simulations, the ratio of the $L_c$-like chains to the total amount of chains ($L_e$ or $L_c$) in each bin was calculated, and considered as the fraction of the $L_c$ phase at the corresponding distance from the protein.

**Lipid–Protein Contacts** The lipid–protein contacts were determined by calculating the number of lipids of specific type within 0.3 nm of the protein with the GROMACS tool `gmx select`. The calculation was done every 1 ns and the contacts were averaged over the repeats. The trajectories were then divided into sections based on the previously defined surface pressure regimes (0–15, 15–30, 30–45, >45 mN/m) and the contacts were averaged over these regimes. The results were normalized by the mole fractions of the lipids and the contact fraction was calculated as the number of contacts of a lipid type with the protein divided by the total number of lipid contacts. The residue specific lipid–protein contacts were determined by calculating the number of lipids of specific type within 0.3 nm of a specific residue.

The SP–PG contacts were mapped onto the protein structure (BETA field in a PDB file), and rendered using VMD to visualize the spatial distribution of the favourable contacts.

#### Additional Results

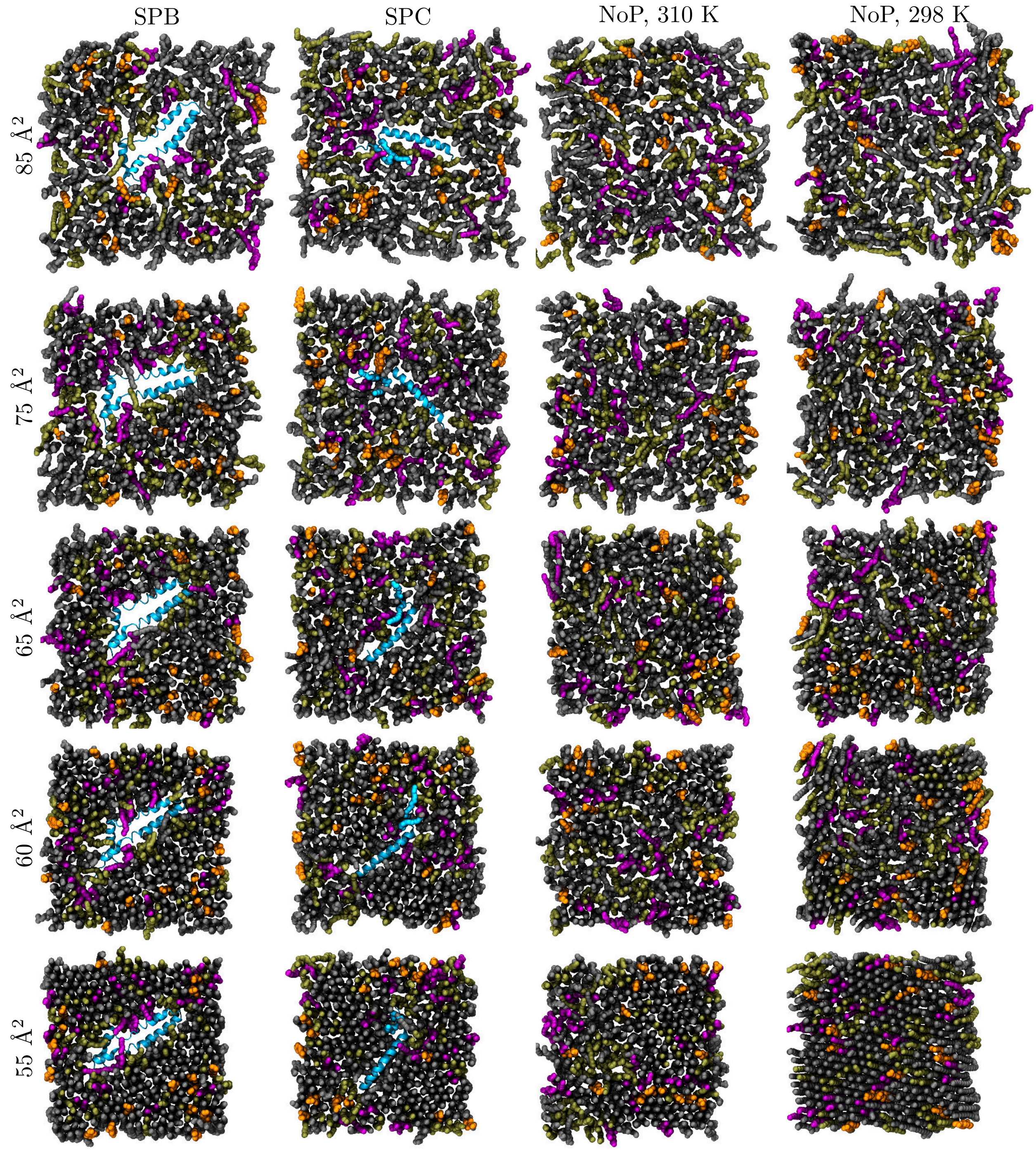


Figure S1: Snapshots from simulations with SP-B, SP-C, and NoP, respectively. The SP-B and SP-C were simulated at 310 K.

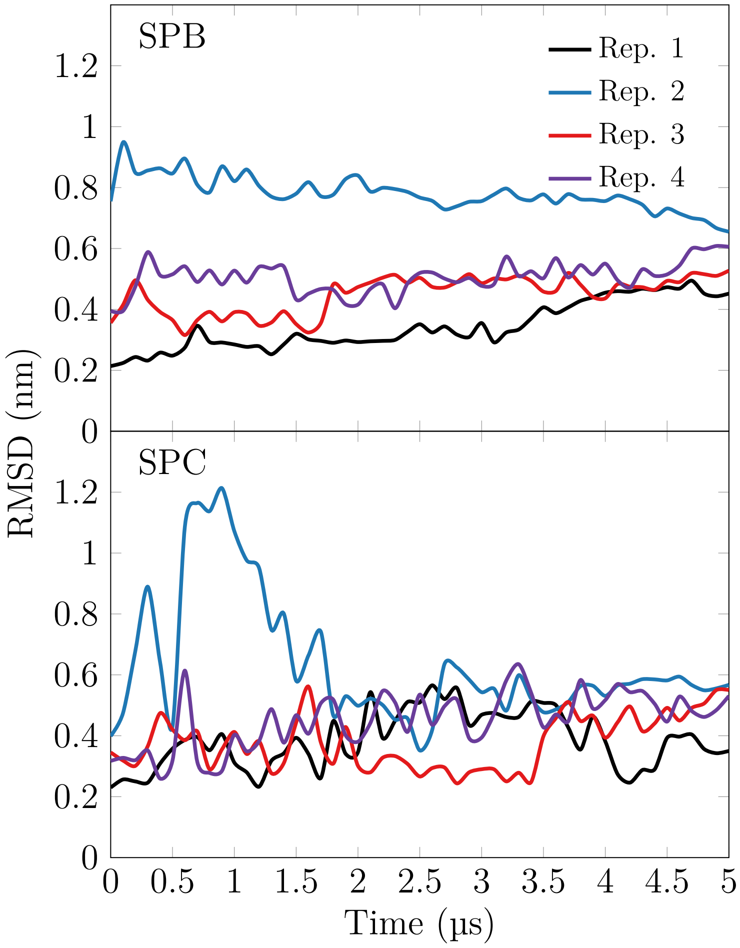


Figure S2: Root mean squared deviation of the protein structures.

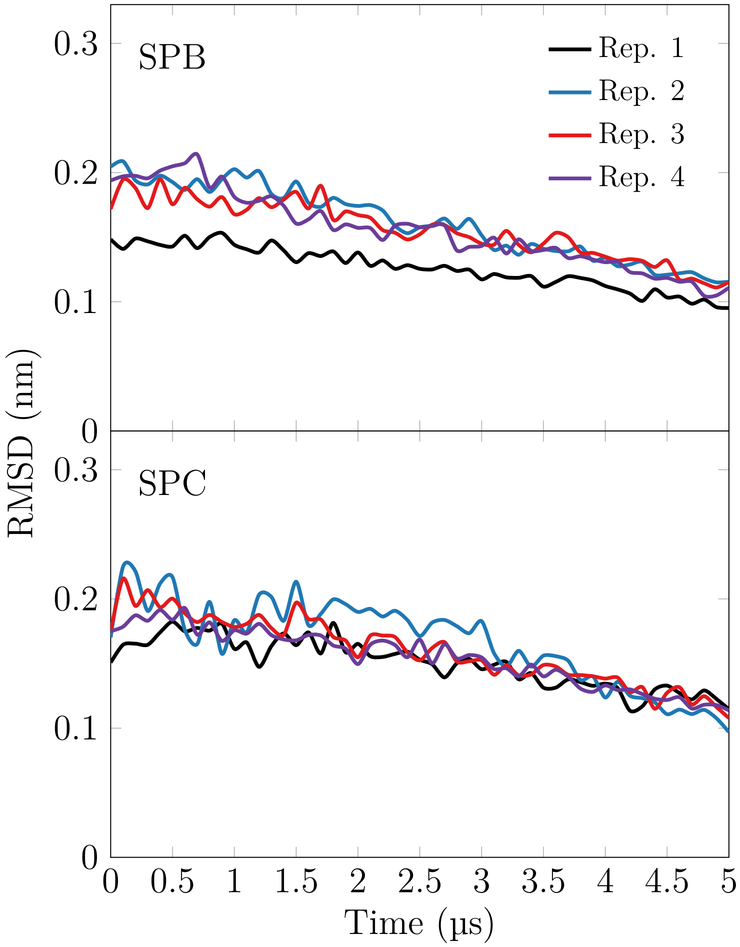


Figure S3: Root mean squared deviation of the protein structures with respect to the previous conformation in the simulation.

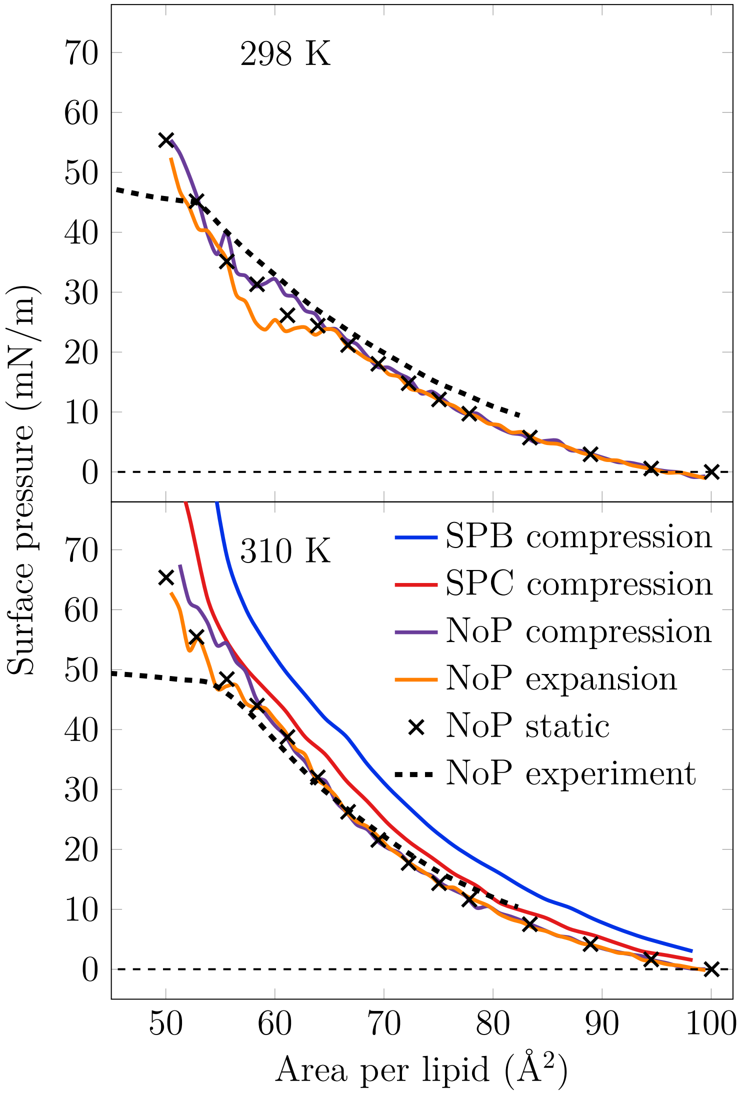


Figure S4: Surface pressure–area isotherms for the monolayer without any proteins at 298 K, and monolayer with SP-B, SP-C, and without any proteins at 310 K.

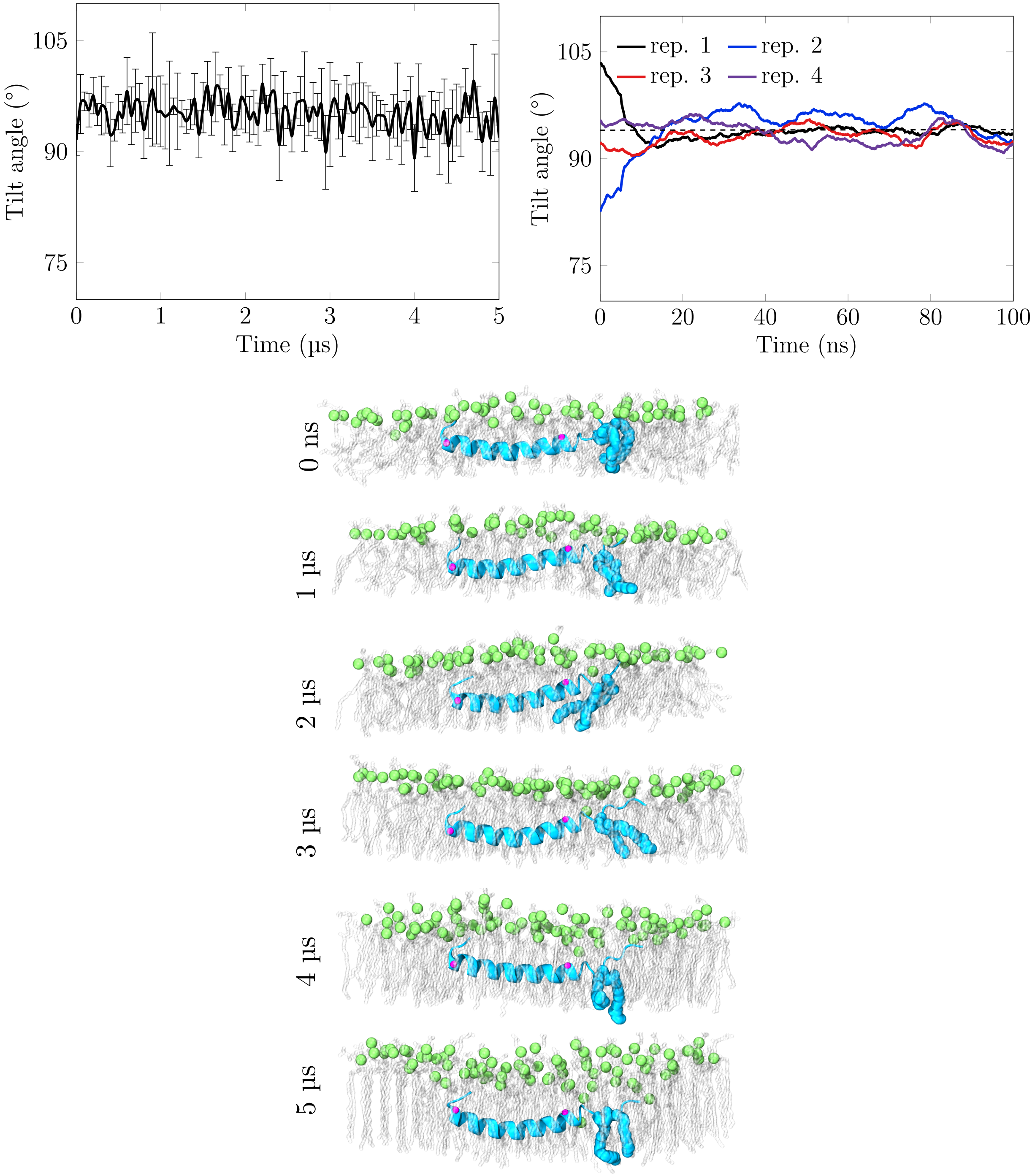


Figure S5: Tilt angle of SP-C in the monolayer in the simulations. On the top left plot, the data is calculated as an average of four repetitions, error estimate as standard error. On the top right plot, the tilt angle of SP-C after placing it parallel or perpendicular to the monolayer plane at constant APL. The bottom snaphots show how SP-C remains close to parallel to the monolayer plane through the simulation. SP-C is shown in blue, phospholipid phosphate atoms in green, lipid tails in gray, and the C-alpha atoms of residues Lys11 and Leu32 are shown in purple.

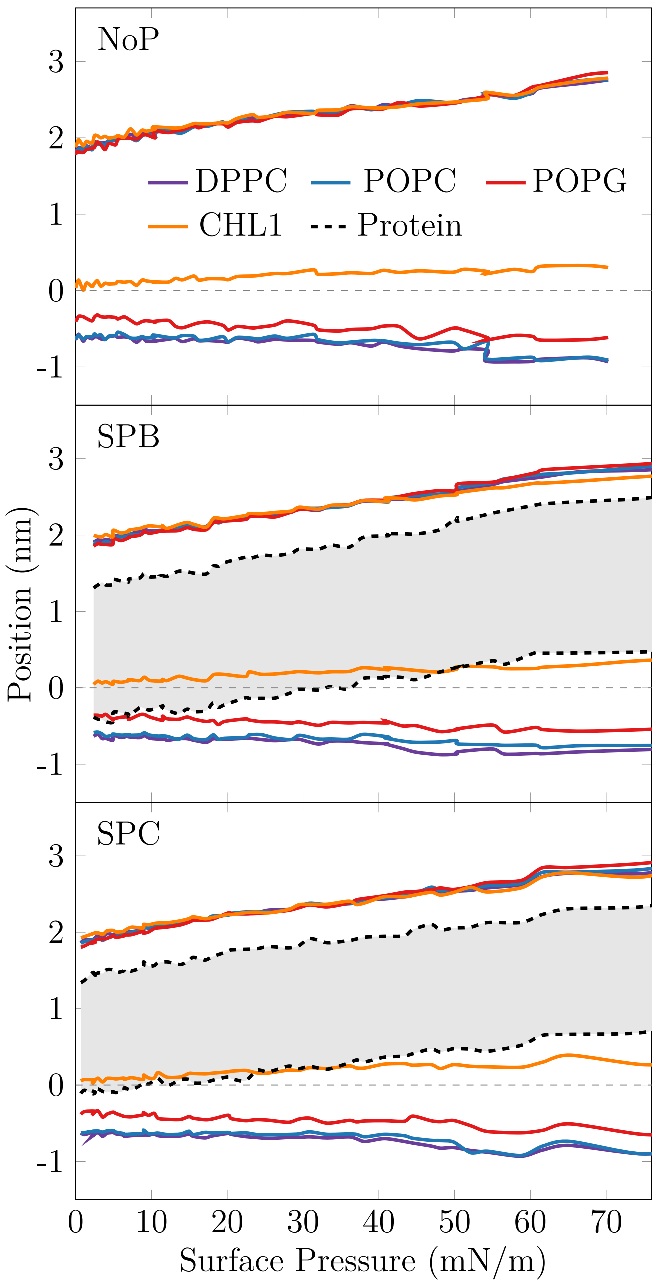


Figure S6: Thickness of monolayers and position of the proteins in the monolayers. The shaded area between the dashed black lines show the position of SP-B and SP-C, respectively. The zero position is defined as the position of the phosphorous atoms of the phospholipids.

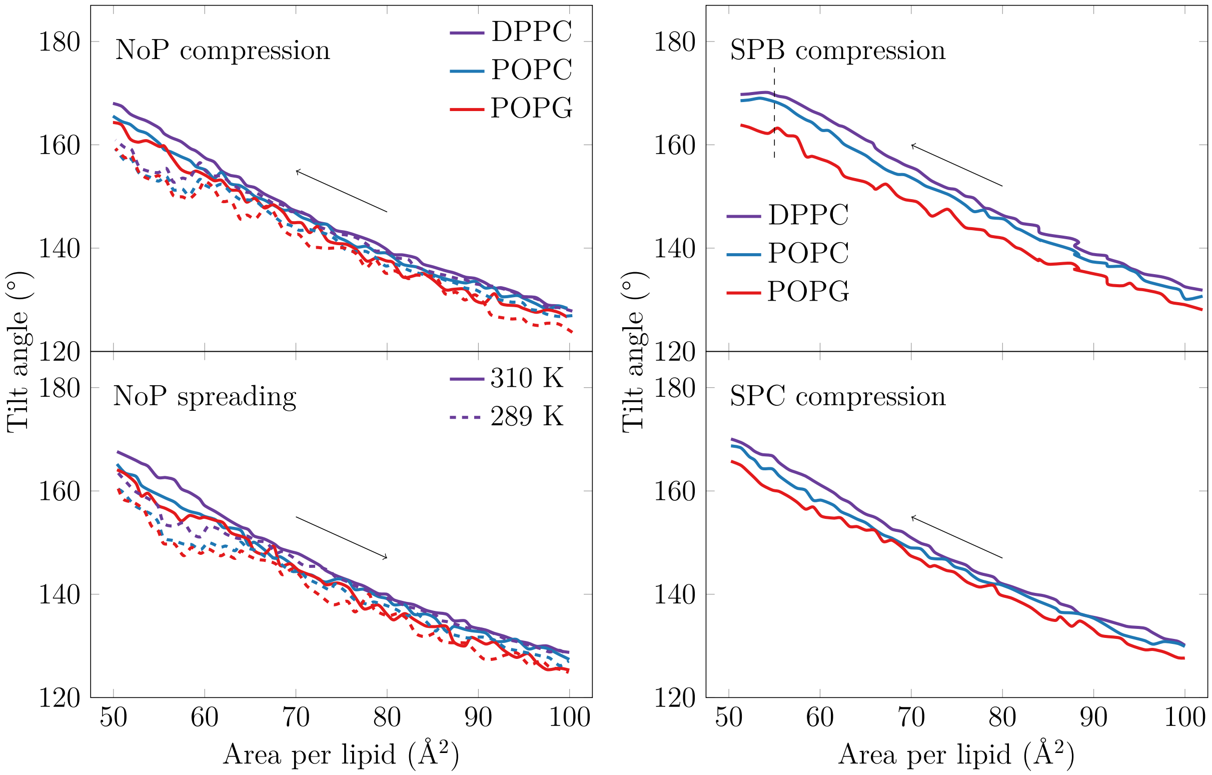


Figure S7: Tilt of the phospholipid acyl chains as a function of compression (APL) and lipid type.

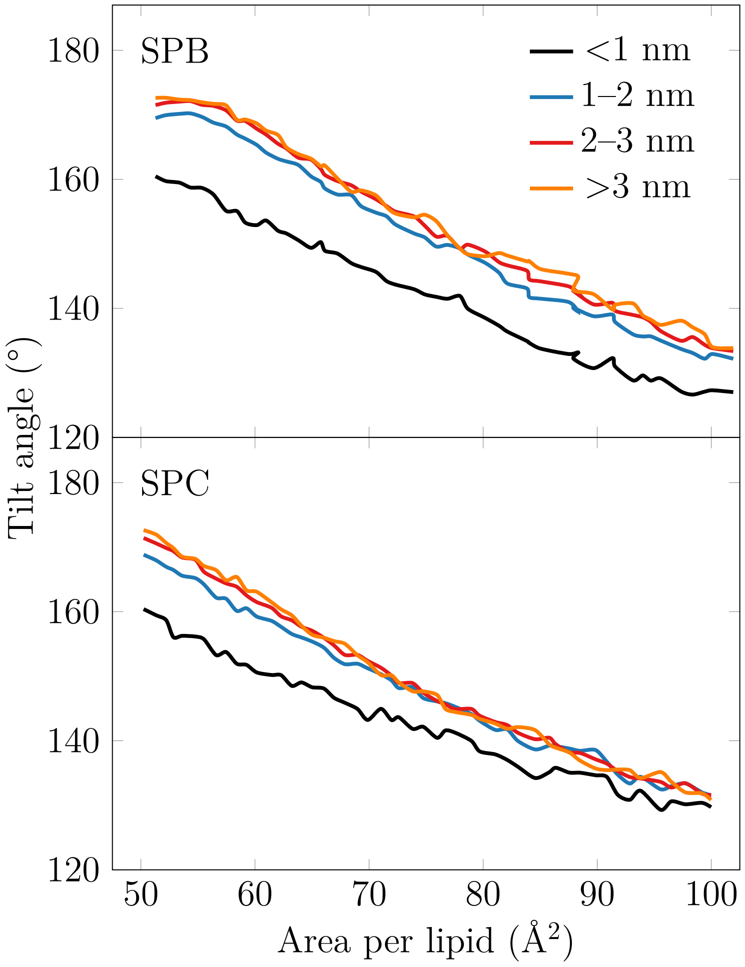


Figure S8: Tilt of the phospholipid acyl chains as a function of compression (APL) and distance from protein.

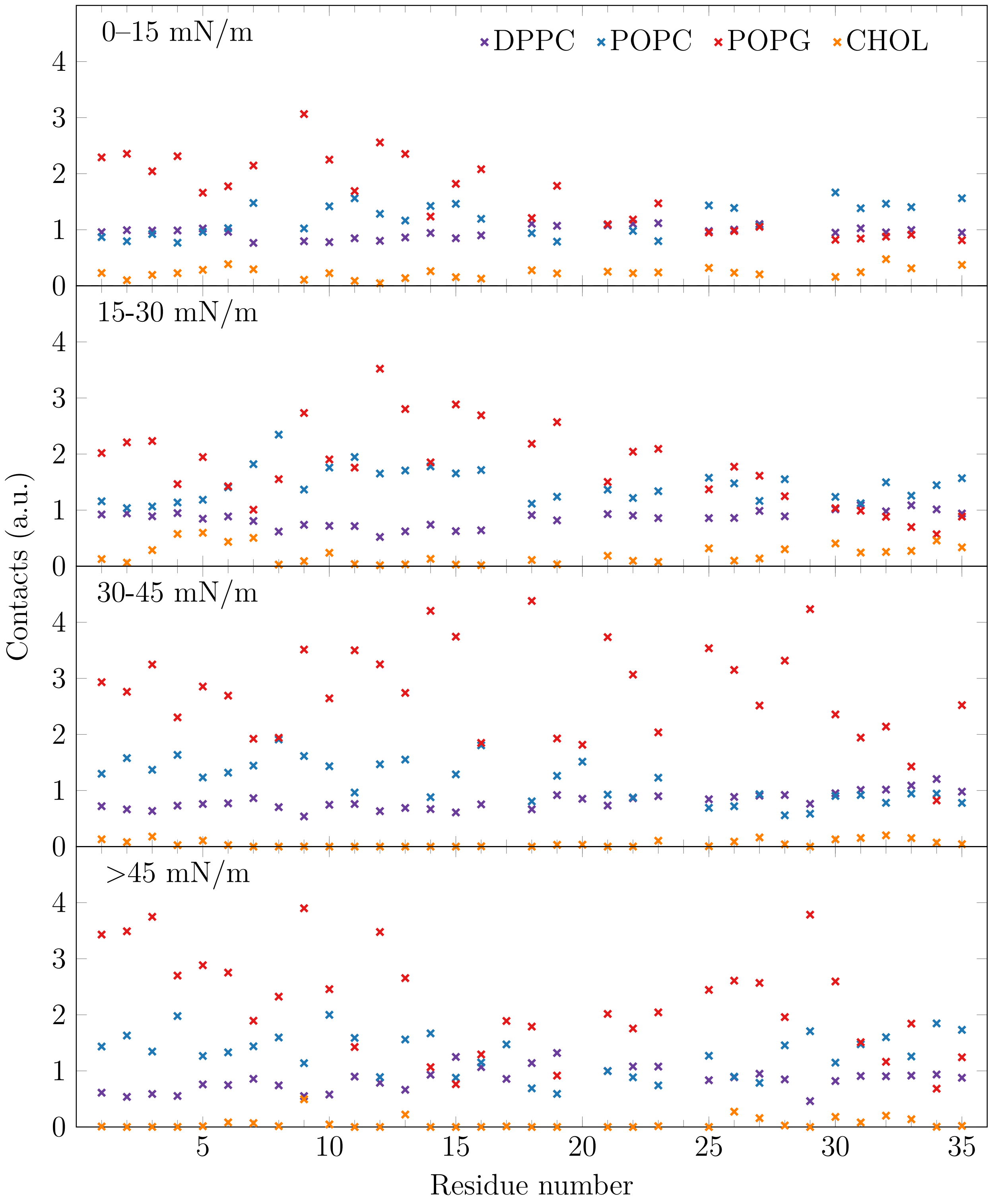


Figure S9: Interactions of SP-C with different lipid types in the different surface pressure intervals.

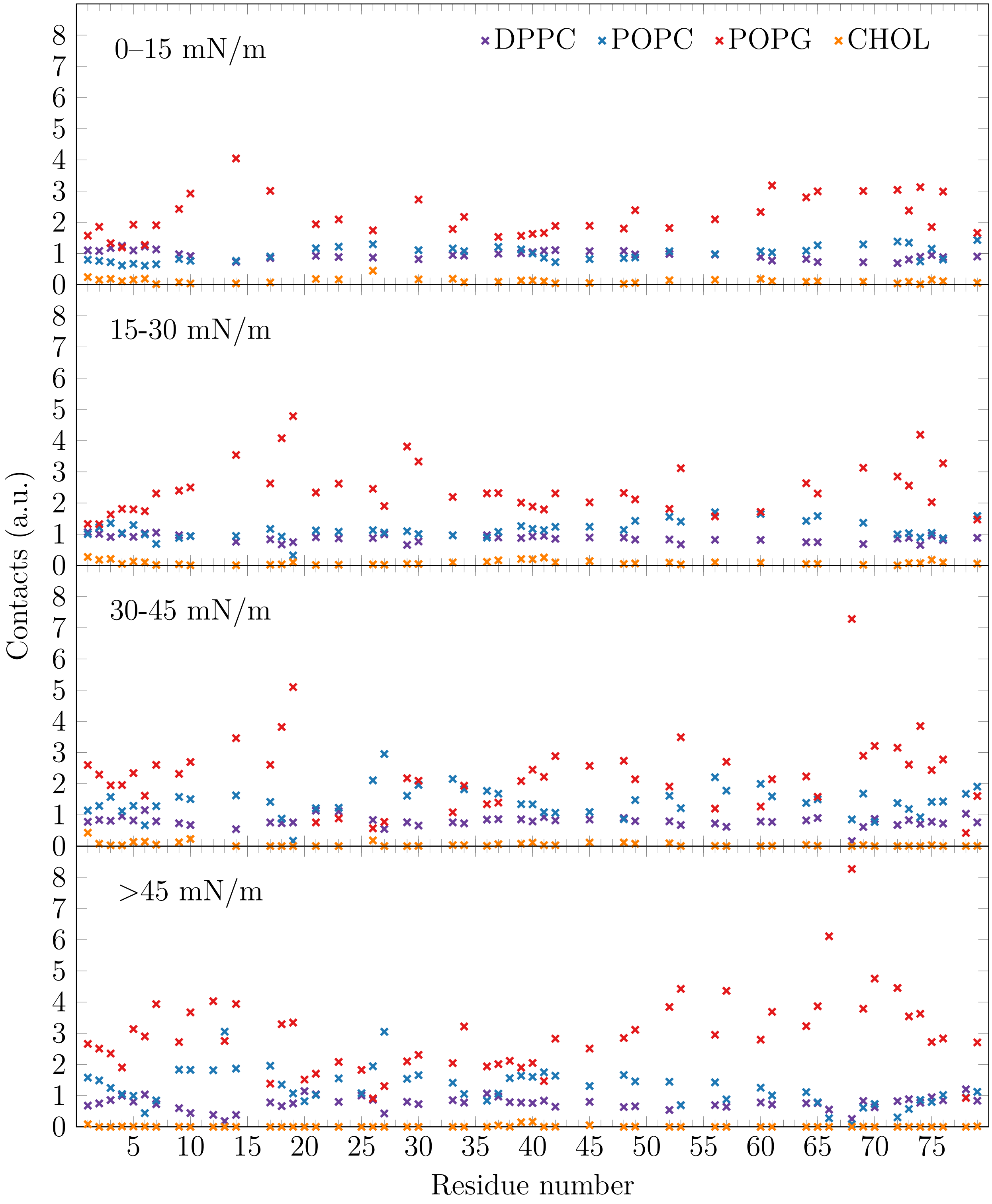


Figure S10: Interactions of SP-B with different lipid types in the different surface pressure intervals.

### Availability of Simulation Data

Simulation data of the NoP, SPB, and SPC monolayers together with the related files are available online at Ref. 6.
